# Effects of electroconvulsive shock on the function, circuitry, and transcriptome of dentate gyrus granule neurons

**DOI:** 10.1101/2024.03.01.583011

**Authors:** Adrienne N. Santiago, Phi Nguyen, Julia Castello-Saval, Hannah M. Chung, Victor M. Luna, Rene Hen, Wei-li Chang

## Abstract

Therapeutic use of electroconvulsive shock (ECS) is particularly effective for treatment-resistant depression. Like other more common forms of antidepressant treatment such as SSRIs, ECS has been shown to increase neurogenesis in the hippocampal dentate gyrus of rodent models. Yet the question of how ECS-induced neurogenesis supports improvement of depressive symptoms remains unknown. Here, we show that ECS-induced neurogenesis is necessary to improve depressive-like behavior of mice exposed to chronic corticosterone (Cort). We then use slice electrophysiology to show that optogenetic stimulation of adult-born neurons produces a greater hyperpolarization in mature granule neurons after ECS vs Sham treatment. We identify that this hyperpolarization requires the activation of group II metabotropic glutamate receptors. Consistent with this finding, we observe reduced expression of the immediate early gene cFos in the granule cell layer of ECS vs Sham subjects. Using single nucleus RNA sequencing, we reveal major transcriptomic shifts in granule neurons after treatment with ECS+Cort or fluoxetine+Cort vs Cort alone. We identify a population of immature cells which has greater representation in both ECS+Cort and fluoxetine+Cort treated samples vs Cort alone. We also find global differences in ECS-vs fluoxetine-induced transcriptomic shifts. Together, these findings highlight a critical role for immature granule cells in the antidepressant action of ECS.

## Introduction

In the United States, 21 million adults representing 8.3% of the adult U.S. population suffer at least one major depressive episode in a single year (NIMH reports for 2021). Selective serotonin reuptake inhibitors (SSRIs) remain the first-line therapy for major depression with a response rate of ∼60% and rate of ∼30-40% for full remission ^1, 2^. Roughly a third of patients remain treatment resistant^3^. Therapeutic use of electroconvulsive shock (ECS) is the gold standard for effectiveness and remission of treatment-resistant depression^4–7^. Yet the question of how ECS therapy produces these powerful effects remains unanswered.

Similar to other more common forms of antidepressant treatment such as fluoxetine^8^, ECS has been shown to increase neurogenesis in the hippocampal dentate gyrus (DG) of rodent models. ECS promotes the activation and proliferation of quiescent neural stem cells^9^, as well as the survival and differentiation of immature adult born granule cells (iGCs)^10^. Dendritic complexity, spinogenesis and axonogenesis of iGCs are all likewise increased following ECS^11–13^. iGCs have also been shown to be necessary for the anxiolytic effects of ECS, as pharmacogenetic deletion of iGCs blocks ECS-induced reduction of latency to feed in the novelty suppressed feeding paradigm^12^. In human subjects, structural MRI of patients with depression reveals an increase in DG volume after ECS treatment, as compared to volume before treatment, and the increase in DG volume correlated with symptom improvement^14^. Interestingly, volume increase was observed in the DG but not the CA1 or CA3 regions of the hippocampus. While we cannot attribute volumetric changes solely to the bodies and neuropil of iGCs, this finding does highlight the importance of DG neuroplasticity in ECS response. Further supporting the role of iGCs, postmortem tissue from patients treated with ECS exhibit greater levels of proteins expressed by iGCs, including doublecortin (DCX) and STMN1^15^. Yet the question of how ECS-induced neurogenesis supports the improvement of depressive symptoms remains unknown.

Several properties of iGCs distinguish them from mature granule cells (mGCs) and provide insight as to their unique role in the DG network. Our group^16, 17^ and others^18, 19^ have shown that iGCs are more active than mGCs, yet paradoxically contribute to overall DG inhibition and maintenance of sparse network activity which may be necessary for optimal execution of DG dependent behaviors^19, 20^. iGCs indirectly suppress mGC activity via inhibitory interneurons^17, 21–24^ and directly inhibit mGCs via group II metabotropic glutamate receptor (mGluRII)-mediated activation of G-protein coupled inwardly rectifying potassium channels ^25^. Interestingly, systemic application of both mGluRII agonists^26, 27^ and antagonists^28^ are associated with anxiolytic effects in mice. This is likely due to the heterogenous actions of mGluRII in different brain regions and cellular compartments. Delivery of systemic mGluRII agonist also reduces expression of the immediate early gene cFos in the DG^26^. Here, we ask whether ECS likewise reduces mGC activity and whether mGluRII-mediated iGC inhibition of mGCs is greater after ECS.

We have used single nucleus RNA sequencing (RNAseq) to profile transcriptomic changes in response to ECS or fluoxetine in a cell-type specific manner. We show here that ECS and fluoxetine not only increase the population of iGCs in keeping with previous reports^11, 12, 29^, but also result in massive but distinct transcriptional changes in mGCs.

## Methods

### Mice

All procedures were conducted in accordance with the U.S. NIH Guide for the Care and Use of Laboratory Animals and the Institutional Animal Care and Use Committees of New York State Psychiatric Institute and Columbia University. Adult male mice were housed in a vivarium grouped 2-4 mice/cage, maintained on a 12-hour light cycle. For all experiments, mice began experimental procedures at 10 weeks of age. C57/6J mice were obtained from Jackson Laboratory. Nestin-CreER^T^^2^ transgenic mice^30^ were bred on a C57/6J background and bred in house (Fig 3).

### ECS

Mice were briefly anesthetized with 2% isoflurane, delivered through a nose cone at an oxygen flow rate of 1 L/min. Electroshocks were delivered to anesthetized mice through ear clipped electrodes at 120 Hz and 50 mA, for a duration of 1 second (Fig 1b). Ringers salt solution was applied to the ear prior to shock to improve conductance. Mice received ECS treatment once every other day for a total of 10 sessions (20 days). Sham mice received isoflurane anesthesia and application of Ringers solution and ear clips, with no delivery of electroshock current. Schematic diagrams illustrating ECS (Fig 1b) and RNAseq (Fig 4a) methods were created using BioRender (2024).

**Figure 1.**
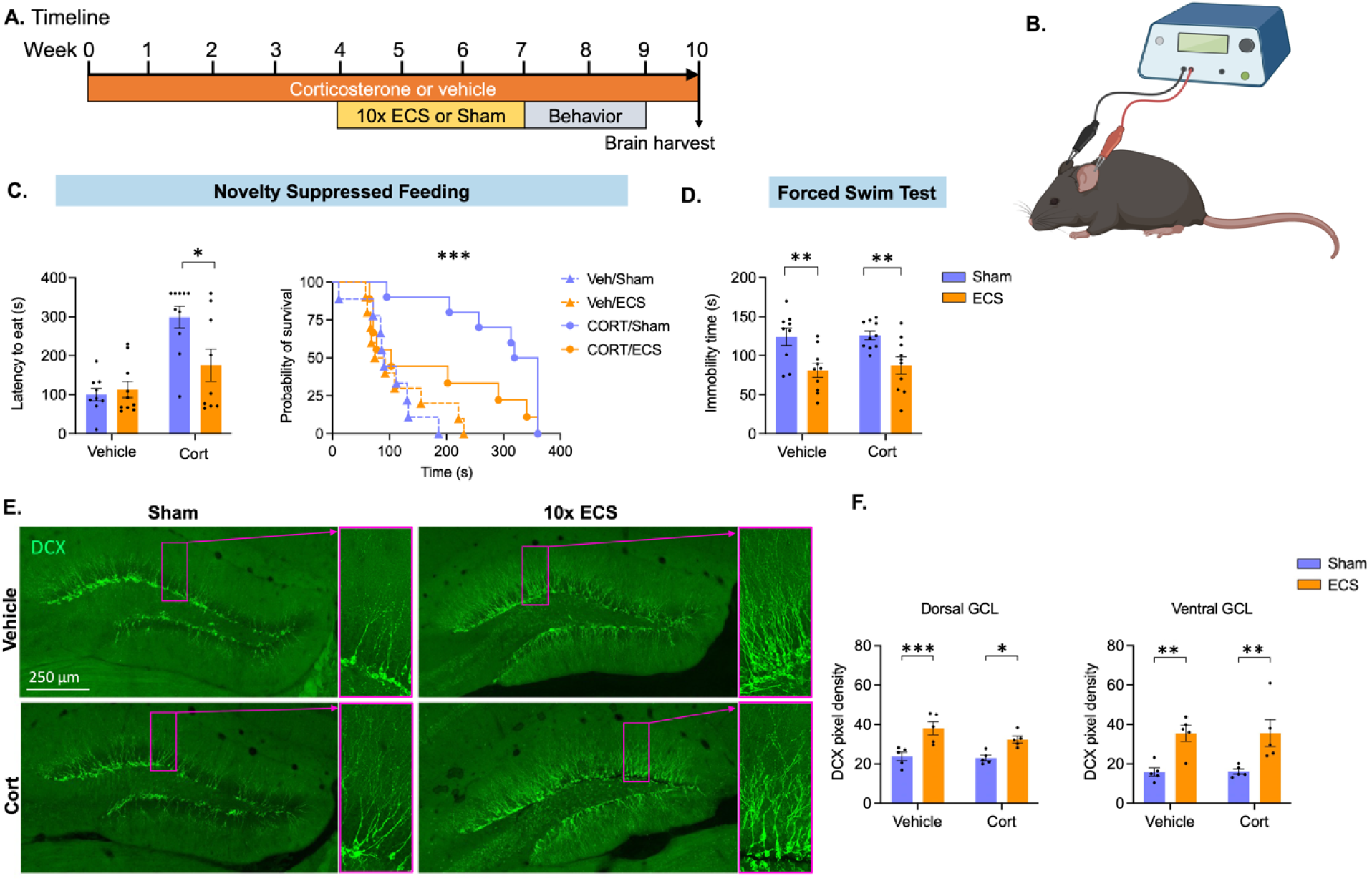
Mouse model of ECS. **A)** Timeline of the experiment. 10-week-old mice were treated with Vehicle or ∼5mg/kg corticosterone (Cort) in drinking water to maintain stress throughout the experiment. After 4 weeks, mice received ECS or Sham treatment every other day for a total of 10 sessions and were tested in paradigms of anxiety-like and depression-like behavior starting 24 hours after the last ECS session. Brains were harvested 1 week after the last behavioral test. **B)** Electroshocks were delivered under isoflurane anesthesia by a Ugo-Basile ECS unit through ear clipped electrodes at 120 Hz, 50 mA for 1s, while Sham mice received anesthesia and ear clip placement, but no electroshock. **C)** In the novelty suppressed feeding test, ECS-treated animals exhibit decreased latency to feed in the Cort group but not in the vehicle group. **D)** In the forced swim test, time spent immobile is reduced after ECS both in the vehicle and Cort groups. **E-F)** Expression of the immature cell marker doublecortin (DCX) is greater in ECS vs Sham subjects. Statistical analysis was performed using two-way ANOVA for normally distributed data (D,F), and the log-rank Mantel-Cox test to report latency to feed (C); (* p ≤ .05, ** p ≤ .01, *** p ≤ .001).

### Drug administration

As in previous publications^31, 32^, Cort (Sigma) was dissolved in .45% betacyclodextrin (vehicle) and delivered via drinking water in opaque bottles replaced every 3-4 days. Mice received 35 ug/ml of Cort, equivalent to 5mg/kg/day. Due to sex differences in response to chronic Cort stress^33^, we limited this study to male mice.

### Focal X-irradiation

Hippocampal irradiation was conducted to eliminate DG neurogenesis, as previously described^8, 25, 34, 35^. After anesthesia with 6 mg/kg sodium pentobarbital i.p., 10-week-old mice were placed in a stereotaxic frame and covered by a lead shield with a 3.22 x 11-mm window positioned to allow focal application of X-irradiation to the hippocampus. X-rays were filtered using a 2 mm Al filter, the corrected dose rate was approximately 1.8 Gy per min and the source to skin distance was 30 cm. A cumulative dose of 5 Gy was given over the course of 2 minutes and 47 seconds over 3 days, with 3 days between each X-ray session. Control mice did not receive radiation, but were anesthetized with irradiated mice throughout the experiment.

### Behavior

Novelty-suppressed feeding (NSF) was performed as previously described (Samuels and Hen 2011). Briefly, mice are food deprived for 16 hrs before placement in a brightly-lit arena with a food pellet taped to the center of the arena. This provides a conflict anxiety test whereby the motivation to eat competes with the risk to enter the center of a brightly lit arena. Forced-swim test was performed as previously described^36^. Briefly, 3L beakers were filled with 2500 ml of tap water at 25C. Time spent actively swimming vs passively floating was recorded using Videotrack software (ViewPoint Behavior Technologies).

### Electrophysiology

We performed whole-cell current clamp recordings on GCs while optogenetically manipulating immature iGCs as previously described^25^.

### IHC

For histology, mice were transcardially perfused with 4% paraformaldehyde (PFA; EMS) in 1X phosphate buffer solution (PBS), after which brains were removed and post-fixed in 4% PFA for 24 h. They were transferred to a 30% sucrose solution in 1X PBS for 2 days, after which they were flash-frozen in methylbutane and coronally sliced on a cryostat (Leica CM 3050S) at a thickness of 40 μm. Immunohistochemistry for DCX and cFos were performed as previously described^17^. Briefly, for DCX, antigen retrieval was performed with heat treatment to 80C in pH8 citrate buffer. Tissue was then incubated in blocking buffer containing 10% normal donkey serum and 0.2% triton for 1 hr. Primary antibody (Cell Signaling Tech #46042S) was diluted 1:400 and incubated with tissue overnight at 4C. After washing 3X in PBS, Alexa Fluor donkey anti-rabbit 488 secondary antibody was used. For cFos, (Cell Signaling Tech #2250), antibody was diluted 1:400 and no antigen retrieval step was applied. All slides were imaged on a Leica SP8 and all IHC images were analyzed with ImageJ software (NIH).

### Flow cytometry for RNAseq

Animals were perfused transcardially with ice-cold 1X PBS and hippocampi were isolated under a dissecting microscope where meninges and excess myelin were removed. For sorting neuronal nuclei for downstream RNA-sequencing, cells were isolated as described^37^. Briefly, hippocampi were mechanically dissociated using a glass tissue homogenizer in isolation medium (HBSS, 15 mM HEPES, 0.6% glucose, 1 mM EDTA pH 8.0). Cells were filtered and then pelleted at 300 g for 5 minutes at 4°C before being resuspended in 22% Percoll (GE Healthcare) and centrifuged at 900 g for 20 minutes with acceleration set to 4 and deceleration set to 1 in order to remove cellular debris. Pelleted nuclei were then washed with isolation medium and incubated in block solution consisting of 1% BSA (Sigma-Aldrich, A2153). Alexa Fluor 488 mouse anti-NeuN (1:1500; Millipore, MAB377X) and DAPI (Sigma-Aldrich, D9542) were used to detect neuronal nuclei. Cell multiplexing oligos (10x Genomics, 3’ CellPlex Kit Set A, 1000261) were incubated with each sample for 5 min at room temperature following washes of primary antibody stain. Gating was based on fluorescence minus one (FMO) negative staining controls. All data analysis was performed using FlowJo™ software. Cells were sorted on a Sony MA900 cell sorter at the Columbia Stem Cell Initiative Flow Cytometry core facility at Columbia University Irving Medical Center. Neuronal nuclei were gated based on forward/side scatter, DAPI+, and NeuN+.

### Single nuclei RNA-sequencing

FACS purified neuronal nuclei were sequenced using the 10 Genomics Single Cell Gene Expression 3’ platform (v3). Nuclei barcoded with cell multiplexing oligos were pooled into 4 samples with each sample containing nuclei from one mouse from each condition (Vehicle, Cort, Fluoxetine + Cort, ECS + Cort). Approximately 15,000 nuclei were loaded into each well of Chromium Chip A according to the manufacturer instructions and combined into droplets with barcoded beads using the Chromium controller. Libraries were prepared by the JP Sulzberger Columbia Genome Center following the instructions in the Chromium Single Cell 3′ Reagent Kits version 3 user guide. The samples were sequenced to an average depth of 40,000-60,000 reads on an Illumina Novaseq 6000 sequencer.

### Single nuclei RNAseq data analysis

Sequenced samples were processed by the JP Sulzberger Columbia Genome Center using the Cell Ranger 7.01 pipeline and aligned to the mm10-2020-A mouse reference transcriptome.

Clustering and differential expression analysis were conducted using Seurat version 4^38, 39^. Cells with fewer than 500 detected genes/cell or over 3000 genes/cell, comprising the lower and upper 5% of cells respectively, or more than 5% mitochondrial RNA were excluded during quality control steps before identification of variable genes in the dataset, cell centering, and scaling. Additionally, 3000 genes were eliminated that were expressed by fewer than 5 cells in the dataset. The “SCTransform” function in Seurat was used for normalization and variance stabilization of molecular count data. Approximately 3000 of the most variable genes were identified and used for calculating nearest neighbors, PCA, and UMAP plots. 50 principal components were calculated using these genes and the first 30 PCs were used for subsequent calculations. Clustering and UMAP plots were computed using Seurat^38, 39^.

Gene lists for mature or immature neuronal profiles were derived from Hochgerner et al., 2018. The “AddModuleScore” function in Seurat was used to generate module scores of mature and immature neuronal profiles. Differentially expressed genes were identified using the “FindMarkers” function in Seurat. Only genes expressed in 25% of the cells in a given condition and a minimal log fold change threshold of 0.25 were included in the differentially expressed gene list. GO analysis was conducted using the Metascape webpage (https://www.metascape.org).

### Statistical analysis

All statistical analyses were preformed using Prism (GraphPad). Generally, normally distributed data were analyzed by two-way ANOVA to assess ECS effects alongside experimental manipulations. Because NSF recordings are capped at 360s, we frequently observed latency to feed measures that were censored at this limit. We therefore used pair-wise Log-rank Mantel-Cox tests to compare these data.

## Results

### Mouse model of ECS

10-week-old C57/Bl6 mice were treated with vehicle or ∼5mg/kg corticosterone (Cort) in drinking water to maintain a stress-like state throughout the experiment, (protocol as published^31^; Fig 1a). After 4 weeks, mice from Vehicle and Cort groups underwent ECS or Sham treatment. To induce ECS, electroshocks were delivered to anesthetized mice through ear clipped electrodes at 120 Hz and 50 mA, for a duration of 1 second (Fig 1b). This current induced tonic/clonic seizures, identifiable by acute arching of the back followed by rhythmic movement of limbs and sustained tension in the tail. Because repeated ECS is more effective than a single dose and more accurately mimics designs for treatment in humans^40–42^, we repeated ECS on alternate days for 10 sessions. Sham mice received 10 sessions of anesthesia and placement of ear clips, but no administration of electroshock. Duration of seizure was unaffected by Cort vs vehicle administration (two-way ANOVA; F(1,9) = 0.07; p = 0.79; Fig S1a).

We first established replication of published behavioral effects of ECS on anxiety-like and depression-like behaviors^12, 40^. During acclimation to behavioral testing, locomotion was observed in a simple arena for 10 min. Total locomotion did not differ significantly between groups (one-way ANOVA: F (3, 35) = 1.378; p=.2655; Fig S1b). For the novelty suppressed feeding test (NSF), mice were deprived of food for 24 hrs before they were given access to a food pellet in the center of a novel arena. Latency to consume food is recorded for up to 360s, before the test is repeated in the home cage. Latency to approach food specifically in the novel arena is an established indicator of anxiety-like behavior^8, 43^. In the novel arena, ECS-treated mice exhibit a decreased latency to feed in the Cort group (p = .0270) that is absent in the vehicle group (p = .6790, Log-rank Mantel-Cox test; Fig 1c). By contrast, ECS had no effect on latency to feed observed within the home cage (two-way ANOVA F (1, 34) = 0.02409; p = 0.8776). In the Forced Swim Test (FST), ECS mice spent less time immobile vs Sham in both the Cort (p= .0107) and Vehicle (p= .0052) treated groups (two-way ANOVA: F (1, 35) = 19.31; p < .0001; Sidak’s test post hoc; Fig 1d). Together, these findings support previous publications establishing the anxiolytic and antidepressant-like effects of repeated ECS, particularly after exposure to chronic stress^12, 40^.

### Effects of ECS on iGCs

A subset of 5 mice per group were analyzed for expression of doublecortin (DCX), which is expressed in the dendrites and somata of immature adult-born granule neurons (Fig 1e). Estimating the relative density of DCX expression in the granule cell layer averaged over the full dorsoventral axis of the DG, we found that ECS treated mice had more DCX expression vs Sham mice, regardless of Cort (p = .0045) vs Vehicle (p = .0005) treatment (Fig 1f; two-way ANOVA: F (1, 16) = 34.57; p < .0001; Sidak’s test post hoc). When we limited our analysis to the dorsal DG, we likewise found that ECS treated mice expressed higher density of DCX vs Sham in both Cort (p=.0185) and vehicle treated mice (p = .0008; two-way ANOVA: F (1, 16) = 27.56; p < .0001; Sidak’s test post hoc). When we limited our analysis to the ventral DG, we again found that ECS treated mice expressed higher density of DCX vs Sham in both Cort (p=.0084) and vehicle treated mice (p = .0080; two-way ANOVA: F (1, 16) = 22.38; Sidak’s test post hoc).

These findings are in keeping with past publications that have reported increased neurogenesis after ECS as well as longer and more complex dendritic branches in DCX immunoreactive dendrites^10, 13^.

### Effects of x-irradiation to ablate DG neurogenesis on ECS efficacy

To assess the role of iGCs in mediating the effects of ECS, we performed focal DG X-irradiation (X-IR) vs Sham (X-sham) followed by ECS vs Sham (E-sham) in Cort treated mice (Fig 2a). X-IR eliminated expression of DCX (Fig 2b), as has been shown previously^8, 25, 34, 35^.

**Figure 2.**
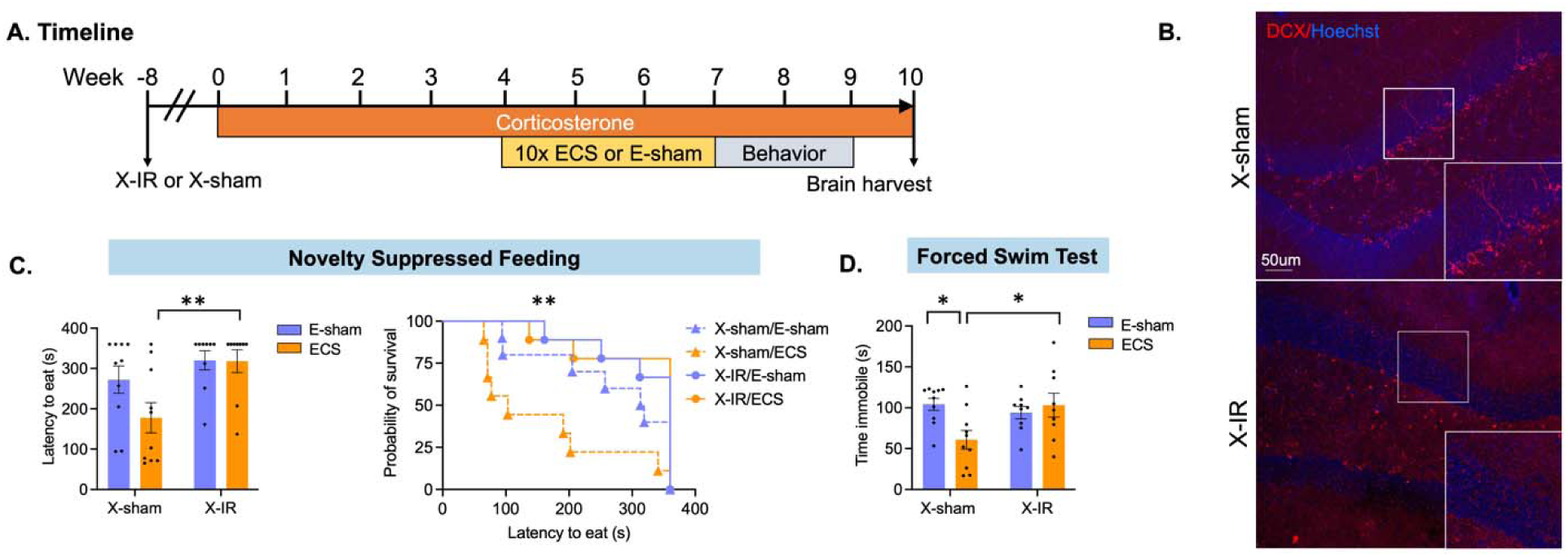
iGCs are required for the behavioral effects of ECS. **A)** Experimental timeline. Mice received X-irradiation targeting the hippocampus (X-IR) or sham (X-sham). After 8 weeks of recovery, all mice received Cort in drinking water for the duration of the experiment. After 4 weeks, mice received 10 sessions of ECS or Sham (E-sham) and were behaviorally tested in the novelty suppressed feeding test and the forced swim test. **B**) Doublecortin (DCX) immunohistochemistry reveals successful ablation of iGCs in X-IR mice vs. X-sham. **C)** On the NSF test, latency to feed is reduced in ECS-treated mice with intact neurogenesis but not in X-IR mice treated with ECS. **D)** On the FST, X-IR mice receiving ECS do not show decreased immobility time when compared to Sham-treated mice. Statistical analysis was performed using two-way ANOVA for normally distributed data (D), and the log-rank Mantel-Cox test to report latency to feed (C); (* p ≤ .05, ** p ≤ .01).

In the NSF test, we observed that ECS treated mice had a higher latency to eat after X-IR vs ECS/X-sham mice (p = 0.0023, log-rank mantel-cox test; Fig 2c). By contrast, X-IR had no significant impact on latency to eat within the home cage for either ECS (p = .3500) or E-sham groups (p = .254).

In the FST, ECS mice were immobile for significantly less time in the X-sham group vs X-IR treated mice (p = 0.0155, two-way ANOVA interaction effect: F (1, 34) = 6.135; p < .0184; Sidak’s test post hoc; Fig 2d). Within the X-sham group, ECS mice spent less time immobile vs E-sham mice (p = .0107). Together, these findings support a critical role for iGCs in mediating ECS-induced improvement of both anxiety-like and depressive-like behaviors.

### Effects of 10x ECS on iGC-driven inhibition of mGCs

Several studies from our group^17, 20, 25^ and others^18, 19^ have suggested that iGCs support DG function by decreasing activity in more mature cells. To assess whether DG activity was generally sparser after ECS vs Sham, we quantified expression of the immediate early gene, cFos, after exposure to home cage (Fig 3a). We found that cFos expression was lower in ECS vs Sham treated mice (Student’s T test; p = .0007; Fig 3b).

**Figure 3.**
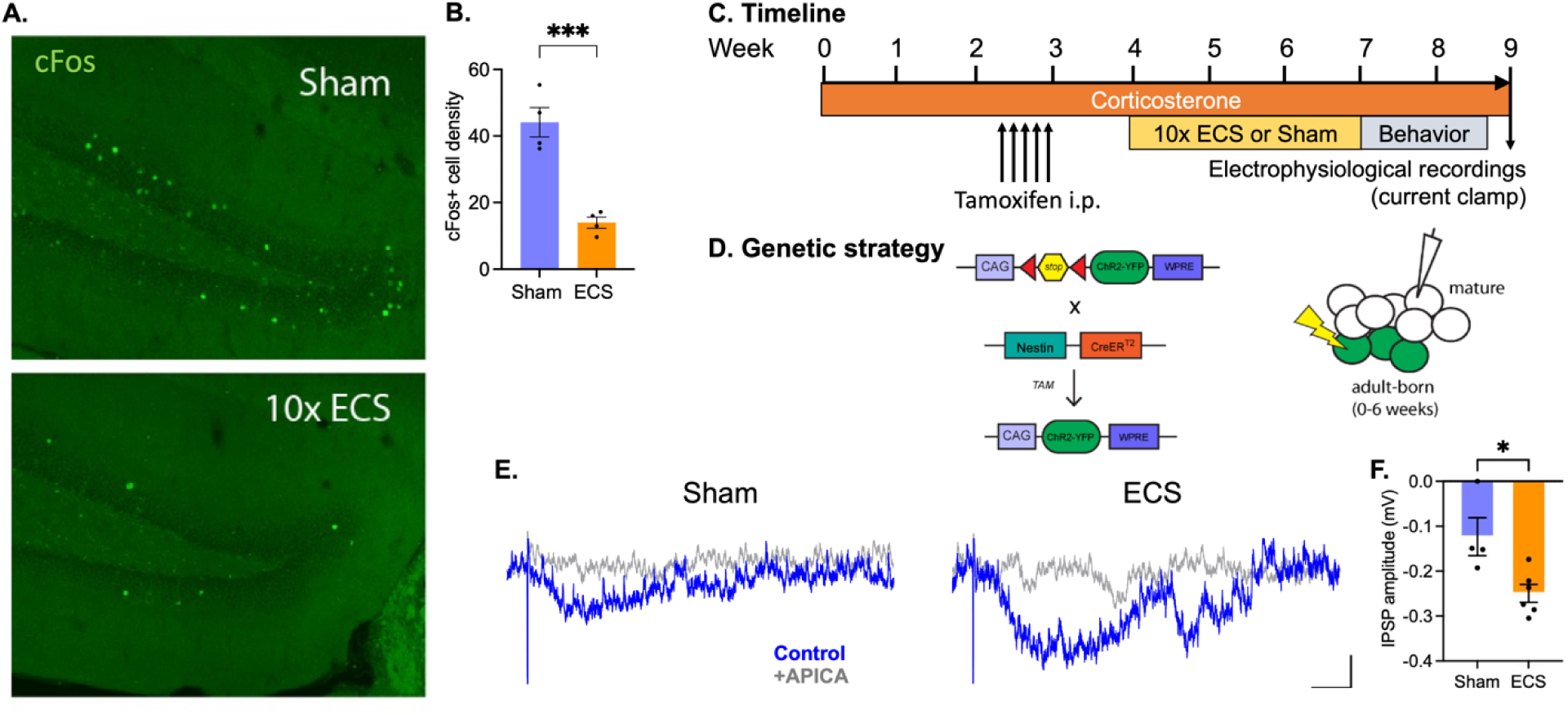
mGluRII mediates iGC-driven inhibition in mGCs after 10x ECS. **A)** cFos expression in the granule cell layer of the dentate gyrus following home cage rest is lower in mice that received 10x ECS vs Sham controls (**B**), suggesting that ECS supports a sparser network at rest. **C)** Experimental timeline. Mice received Cort in drinking water for 4 weeks. Six weeks after tamoxifen injection to induce expression of ChR2-EYFP in iGCs, whole-cell recordings were performed in mature granule cells in DG brain slices. Delivery of 10x ECS or Sham occurred one week after tamoxifen-induced Cre recombination. **D)** Breeding strategy and experimental protocol for electrophysiological approach. Nestin-CreER^T2^ mice were crossed with floxed Channelrhodopsin-EYFP (ChR2-EYFP) expressing mice. Optogenetic stimulation of iGCs evoked responses in mGCs. **E)** We optogenetically stimulated 0-6-week-old iGCs to evoke inhibitory postsynaptic potential in mGCs in the presence of 20μM NBQX, 50μM APV, and 20μM bicuculline (blue trace). These inhibitory signals were blocked by bath application of the mGluRII antagonist APICA (500 μM; grey trace; scale bar = 200 ms, 100 mV). **F)** mGluRII-mediated IPSP had greater negative amplitude after ECS vs Sham in Cort treated mice. (* p ≤ .05, *** p ≤ .001).

In addition to inhibitory cell recruitment, iGCs are capable of directly inhibiting mGCs via mGluRII-mediated postsynaptic hyperpolarization^25^. To determine whether this type of inhibition was enhanced by ECS vs Sham in subjects maintained on Cort, we used whole-cell current clamp recordings in DG brain slices (Fig 3c). For these experiments, we crossed a Nestin-CreER^T^^2^ mouse line with a floxed channelrhodopsin-EYFP (ChR2-EYFP) expressing mouse line, which enabled us to inducibly express ChR2-EYFP in iGCs upon delivery of tamoxifen (Fig 3D). We optogenetically stimulated 0-to 6-week-old iGCs and recorded synaptic responses in the presence of AMPA-, NMDA-, and GABAA-receptor antagonists (Luna et al., 2019) (Fig 3e). We observed a significantly larger inhibitory response in mGCs of ECS-treated mice than Sham controls (Fig 3f; Sham=-0.12±0.04 mV, N=4; ECS=0.25±0.02, N=6; Student T-test, p = .0162). The inhibitory currents were blocked with bath application of the mGluRII antagonist, APICA, indicating that direct iGC hyperpolarization of mGCs requires mGluRII, as previously shown (Luna et al., 2019). Together these findings suggest that iGC activation may contribute to the sparsity of the granule cell network by engaging mGluRII present on mGCs.

### Both ECS and Fluoxetine drive transcriptomic shifts indicating greater neuroplasticity and more prominent immature phenotype among granule neurons

To characterize granule neuron response to antidepressant treatment, we performed single nucleus RNA sequencing on dissected hippocampi from four treatment groups: vehicle (Veh), Cort, Cort plus fluoxetine (Flx), and Cort plus 10x ECS (N=3 per group; Fig 4a). Neuronal nuclei were isolated using fluorescent activated cell sorting (FACS) for NeuN+ nuclei. Transcriptomic profiles of hippocampal neuronal nuclei were generated and granule cells were identified by expression of Prox1, Dock10, and Stxbp6 (N=8,318 nuclei). We then used Uniform Manifold Approximation and Projection (UMAP) to reduce the dimensionality of granule neuron RNA profiles to two axes in order to visualize relative similarity vs difference in expression profile between individual neurons (Fig 4b). We observed that nuclei from Veh, Cort, Flx, and ECS groups clustered more tightly within vs across groups, indicating that granule neurons from each group are transcriptomically distinct (Fig 4b). If Flx or ECS simply reversed the effect of Cort, we may expect to see more overlap between these groups and the vehicle group. Instead, we observe discrete transcriptomic profiles for each group.

**Figure 4.**
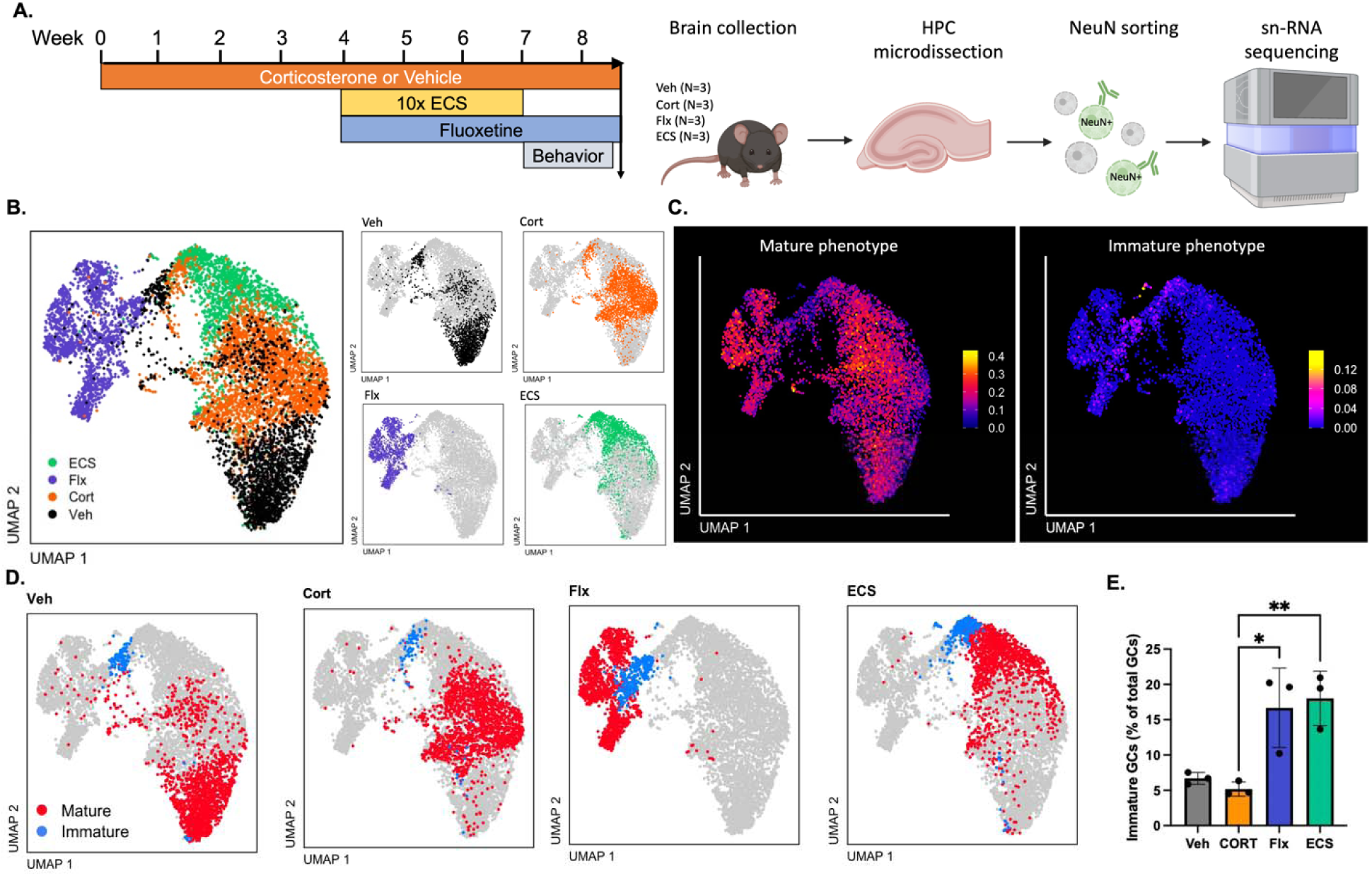
Both ECS and Fluoxetine drive transcriptomic shifts indicating greater neuroplasticity and more prominent immature phenotype among granule neurons. **A**) Single nucleus RNA sequencing was performed on hippocampal nuclei from four treatment groups: vehicle (Veh), corticosterone (Cort), Cort plus fluoxetine (Flx), and Cort plus 10x ECS (N=3 per group). Neuronal nuclei were identified with NeuN antibody followed by cell sorting. UMAP clustering of all neurons revealed a granule neuron population which was used for all subsequent analyses. **B**) UMAP projections of the granule neuron population, labeled by treatment condition. **C**) Genes identified in Hochgerner et al 2018 as those predominantly expressed in either mature or immature cell types were used to produce mature vs immature phenotype module scores. UMAP projections depict granule neurons from all treatment conditions, while color encodes intensity of expression of mature vs immature module scores. **D**) Mature vs immature phenotype module score was used to identify mature vs immature cell clusters. UMAP clustering of mature (red) and immature (blue) nuclei by treatment condition are depicted. **E**) The percent of granule neurons in the immature cell cluster is greater for Flx and ECS treated mice vs Cort alone. Each point represents a single mouse (One-way ANOVA with Tukey posthoc, * p ≤ .05, ** p ≤ .01).

To further investigate the role of immature cells in antidepressant treatment, we used differentially expressed genes (DEGs) identified by Hochgerner et al 2018 as characteristic of immature vs mature cells to create module scores of a mature vs immature phenotype (Fig 4c). We found that, in our sample, mature vs immature phenotype cells produced distinct clusters (Fig 4d). Interestingly, the percent of neurons in the immature phenotype cluster differed by treatment group (Fig 4e; one-way ANOVA, F (3, 8) = 11.00, p = .0044). Post hoc Tukey’s tests revealed that both Flx (p = .0152) and ECS groups (p = .0083) had a significantly greater proportion of immature-clustered granule cells vs Cort.

Both ECS and Flx exhibited upregulation (vs Cort) of key genes known to play a critical role in neurodevelopment and neuroplasticity (see Supplemental Table 1 for a full list of DEGs for ECS vs Cort and Flx vs Cort). These include Neuregulin 1 and 3 (Nrg1, Nrg3), Neurexin 3 (Nrxn3), and Ncam2. ECS also upregulates Nrxn1, while Flx upregulates both BDNF and the gene encoding TrkB (Ntrk2). Both ECS and Flx also exhibit downregulation of genes associated with mature granule cell identity^44^, including Calb1, Dock10, and Rbfox3.

Consistent with our UMAP results (Fig 4b), we find strong differences in Flx- and ECS-induced transcriptomic shifts. While Flx induces upregulation of 1125 DEGs and downregulation of 645 DEGs, ECS upregulates only 209 DEGs and downregulates 1560 DEGs (Fig 5a,b). Flx also induces a stronger transcriptomic response, with greater average fold change and -LogP value for DEGs (Fig 5c,d). Thus, ECS achieves equivalent behavioral antidepressant effects vs Flx while unexpectedly inducing a more modest transcriptomic shift.

**Figure 5.**
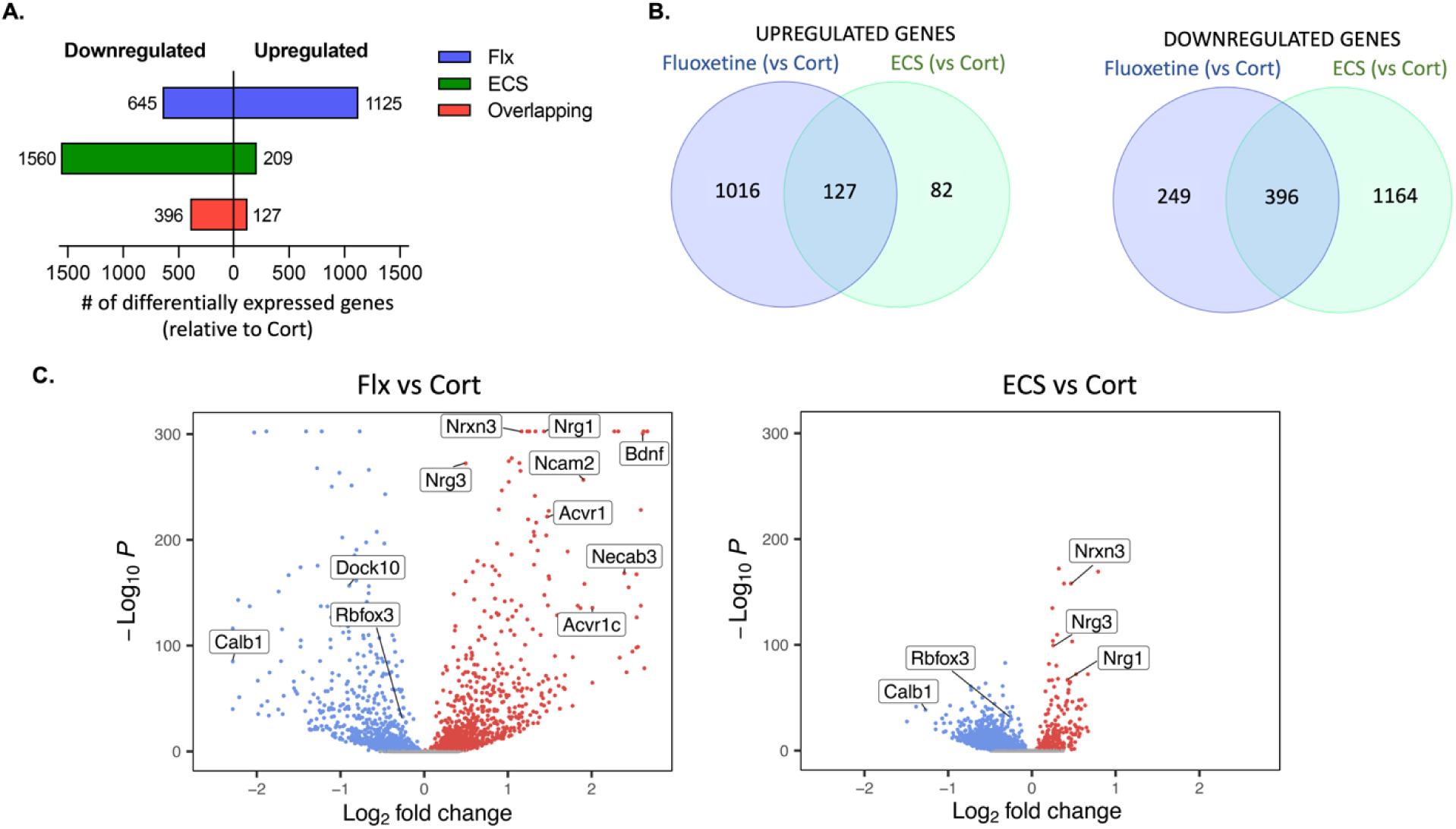
Treatment with fluoxetine induces greater transcriptome upregulation while ECS induces a downward transcriptomic shift. **A)** Representation of the number of differentially expressed genes (DEGs) that are upregulated (right) vs downregulated (left) for Flx and ECS groups, relative to Cort expression. **B**) Venn diagram depicting overlap of upregulated vs downregulated DEGs in Flx and ECS groups, vs Cort. **C**) Volcano plot depicting expression level (-Log of the p value by Log2 fold change) of DEGs for Flx and ECS groups, vs Cort.

## Discussion

### Summary of results

Here, we demonstrated that ECS-induced anxiolytic- and antidepressant-like effects require iGCs. We show for the first time that ECS amplifies direct iGC-induced hyperpolarization of mGCs via mGluRII. We then used single nucleus RNAseq to characterize the transcriptomic shift induced by ECS or Flx vs Cort alone. We show that ECS and Flx both induce a shift towards greater expression of an immature vs mature granule neuron profile, but that the overall expression patterns of the two interventions are distinct.

### iGCs and the sparse DG

In this report, our behavioral, histochemical, electrophysiological, and transcriptomic data converge in their support of the hypothesis that iGCs play a critical role in the antidepressant action of ECS. While several publications have detailed the neurogenic effects of ECS^9, 10^, this is the first study to demonstrate that ECS facilitates iGC-driven inhibition of mGCs. We show here that chronic ECS reduces DG cFos expression over a week after seizure cessation. This finding provides a sharp contrast to the acute effect of ECS, which is known to upregulate activity-dependent genes like cFos post seizure^45^.

Optogenetic activation of iGCs has been shown to increase the sparsity of mGC calcium transients, while optogenetic inhibition of iGCs decreases mGC sparsity^19^. Likewise, a transgenic increase in neurogenesis decreases granule cell activity, while X-IR to ablate neurogenesis increases general granule cell activity^22^. One recent publication found that X-IR did not significantly reduce mGC activity in cue or place cells during contextual discrimination, but instead impaired mGC contextual remapping^35^. This finding suggests that iGCs may have task-dependent effects on mGC activity. We hypothesize that iGC-induced suppression of mature GNs contributes to the sparse DG network activity that has been shown to support stress-resilience^20^.

Our past work has shown that iGCs can directly inhibit mGCs via mGluRII-mediated activation of G protein-coupled inwardly rectifying potassium currents (GIRKs)^25^. Group II mGluRs include both mGluR2 and mGluR3, but only mGluR2 has been shown to engage GIRKs in granule neurons^46^. mGluR2 is expressed on both axon terminals and proximal dendrites of granule cells^46^. While axon terminal mGluR2 has been shown to limit neurotransmitter release via GIRK-independent reduction of calcium conductance^47^, mGluR2 expression on proximal dendrites of mGCs has been shown to engage GIRK-induced hyperpolarization in a dendrite-specific manner^46^.

Interestingly, expression of both mGluR2 and GIRK is absent in very immature granule cells and increases with maturity^48, 49^. This suggests that the electrophysiological effects of mGluR2 that we observe are specific to the more mature mGC population. Interestingly the antidepressant-like effects of fluoxetine are mediated by 5-HT1A receptors (an inhibitory GPCR like mGluR2) also expressed on mGCs^50^. Furthermore, directly inhibiting mGCs (via inhibitory DREADDs) produces a stress resilient phenotype^20^. These convergent results suggest that the inhibition of mGCs is a key component in the response to antidepressants and in resilience to stress^20, 50^.

#### ECS and Flx increase of transcription factors associated with neuroplasticity and immaturity

Given the fact that ECS and fluoxetine both produce antidepressant-like effects and both require neurogenesis, we hypothesized that both manipulations would produce similar transcriptomic changes in the DG. We show here that both Flx and ECS increase transcription of several genes known to play a critical role in iGC survival, growth, migration and connectivity, including the neurexins^51^, neuregulins^52^, and NCAM2^53^ highlighted in this work. We then used the Hochgerner 2018 study to profile granule neurons as having a more immature vs mature phenotype, and found a greater number of immature-phenotype cells in the Flx and ECS populations vs controls. This finding parallels work using BrdU and DCX labeling, in situ hybridization, and neuronal tracing to show that both Flx^31^ and ECS^9, 10^ promote the survival, growth, and dendritic arborization of adult-born iGCs. ECS may additionally lengthen the amount of time adult-born cells retain their immature phenotype, since ECS has also been shown to suppress mature marker expression during the late maturation stage of neurogenesis^10^, as in our RNAseq results (Fig 4 and Fig 5). This dematuration-Iike phenotype^54^ was initially thought to result in an increase in DCX and calretinin expression (two markers of iGCs) as well as a decrease in calbindin (a marker of maturity). However, we recently showed that the expression of DCX and calretinin is exclusively driven by adult-born iGCs^55^. Therefore, it is unlikely that the increase in DCX we observe here is due to dematuration.

There remains the possibility that, in addition to stimulating neurogenesis, ECS and Flx also induce an immature-phenotype among embryonically born mGCs or adult-born granule cells that are well outside the 4-8-week window commonly associated with immaturity. One of the advantages of RNAseq is the ability to profile maturity vs immaturity according to a wide array of gene expression^56, 57^, rather than based on a single expression factor, such as DCX. This allows us to identify a larger, more diverse population of cells with an immature expression profile^56^. This population may very well include dematured cells and/or reflect slower maturation of iGCs. Here, we observe that immature and mature populations are quite distinct in the Cort and Veh UMAPs, but cluster together in the Flx and ECS UMAPs, indicating greater ambiguity between iGC and mGC subclusters in antidepressant-treated granule cells. These data suggest that antidepressant treatment results in a greater number of cells in the transitional space between immature and mature phenotype clusters, which could be facilitated by either increased dematuration of mGCs or slower maturation of iGCs, in addition to the well documented increase in neurogenesis.

#### Conclusions

Overall, these findings provide insight into the neurobiological mechanisms underlying the antidepressant effects of ECS. We found that ECS-induced stress resilience requires adult hippocampal neurogenesis, and that ECS amplifies iGC-induced hyperpolarization of mGCs via mGluRII. Together with previous studies showing that inhibiting the DG can produce antidepressant-like effects, these findings point to potentially novel therapeutic targets to treat depression.

We also found that ECS and Flx both induce a transcriptomic shift towards a more immature phenotype in granule neurons, which is consistent with the fact that adult hippocampal neurogenesis is necessary for some of their behavioral effects^12, 31^. However, despite these similarities, both treatments elicit distinct transcriptomic changes in mature neurons of the DG, which may be related to distinct mechanism of action and possibly to their distinct antidepressant profile in humans.

## Acknowledgements

This work was generously supported by the Hope for Depression Research Foundation (RH), the Jane Coffin Childs Foundation (PN) and the NIMH: R01MH068542 (RH), K08MH122893 (WC), T32MH015144 (AS and PN) and NIA K01 AG054765 (VL). We are grateful to Dr. Joshua Levitz for his insights regarding mGluRII.

## Supplemental methods and figures

## IHC

For histology, mice were transcardially perfused with 4% paraformaldehyde (PFA; EMS) in 1X phosphate buffer solution (PBS), after which brains were removed and post-fixed in 4% PFA for 24 h. They were transferred to a 30% sucrose solution in 1X PBS for 2 days, after which they were flash-frozen in methylbutane and coronally sliced on a cryostat (Leica CM 3050S) at a thickness of 40 μm. Immunohistochemistry for DCX and cFos were performed as previously described 17. Briefly, for DCX, antigen retrieval was performed with heat treatment to 80C in pH8 citrate buffer. Tissue was then incubated in blocking buffer containing 10% normal donkey serum and 0.2% triton for 1 hr. Primary antibody (Cell Signaling Tech #46042S) was diluted 1:400 and incubated with tissue overnight at 4C. After washing 3X in PBS, Alexa Fluor donkey anti-rabbit 488 secondary antibody was used. For cFos, (Cell Signaling Tech #2250), antibody was diluted 1:400 and no antigen retrieval step was applied. All slides were imaged on a Leica SP8 and all IHC images were analyzed with ImageJ software (NIH).

### Flow cytometry for RNAseq

Animals were perfused transcardially with ice-cold 1X PBS and hippocampi were isolated under a dissecting microscope where meninges and excess myelin were removed. For sorting neuronal nuclei for downstream RNA-sequencing, cells were isolated as described ^37^. Briefly, hippocampi were mechanically dissociated using a glass tissue homogenizer in isolation medium (HBSS, 15 mM HEPES, 0.6% glucose, 1 mM EDTA pH 8.0). Cells were filtered and then pelleted at 300 g for 5 minutes at 4°C before being resuspended in 22% Percoll (GE Healthcare) and centrifuged at 900 g for 20 minutes with acceleration set to 4 and deceleration set to 1 in order to remove cellular debris. Pelleted nuclei were then washed with isolation medium and incubated in block solution consisting of 1% BSA (Sigma-Aldrich, A2153). Alexa Fluor 488 mouse anti-NeuN (1:1500; Millipore, MAB377X) and DAPI (Sigma-Aldrich, D9542) were used to detect neuronal nuclei. Cell multiplexing oligos (10x Genomics, 3’ CellPlex Kit Set A, 1000261) were incubated with each sample for 5 min at room temperature following washes of primary antibody stain. Gating was based on fluorescence minus one (FMO) negative staining controls. All data analysis was performed using FlowJo™ software. Cells were sorted on a Sony MA900 cell sorter at the Columbia Stem Cell Initiative Flow Cytometry core facility at Columbia University Irving Medical Center. Neuronal nuclei were gated based on forward/side scatter, DAPI+, and NeuN+.

### Single nuclei RNAseq data analysis

Sequenced samples were processed by the JP Sulzberger Columbia Genome Center using the Cell Ranger 7.01 pipeline and aligned to the mm10-2020-A mouse reference transcriptome. Clustering and differential expression analysis were conducted using Seurat version 4 ^38, 39^. Cells with fewer than 500 detected genes/cell or over 3000 genes/cell, comprising the lower and upper 5% of cells respectively, or more than 5% mitochondrial RNA were excluded during quality control steps before identification of variable genes in the dataset, cell centering, and scaling. Additionally, 3000 genes were eliminated that were expressed by fewer than 5 cells in the dataset. The “SCTransform” function in Seurat was used for normalization and variance stabilization of molecular count data. Approximately 3000 of the most variable genes were identified and used for calculating nearest neighbors, PCA, and UMAP plots. 50 principal components were calculated using these genes and the first 30 PCs were used for subsequent calculations. Clustering and UMAP plots were computed using Seurat ^38, 39^.

---

**S1. Supplemental figure 1.**
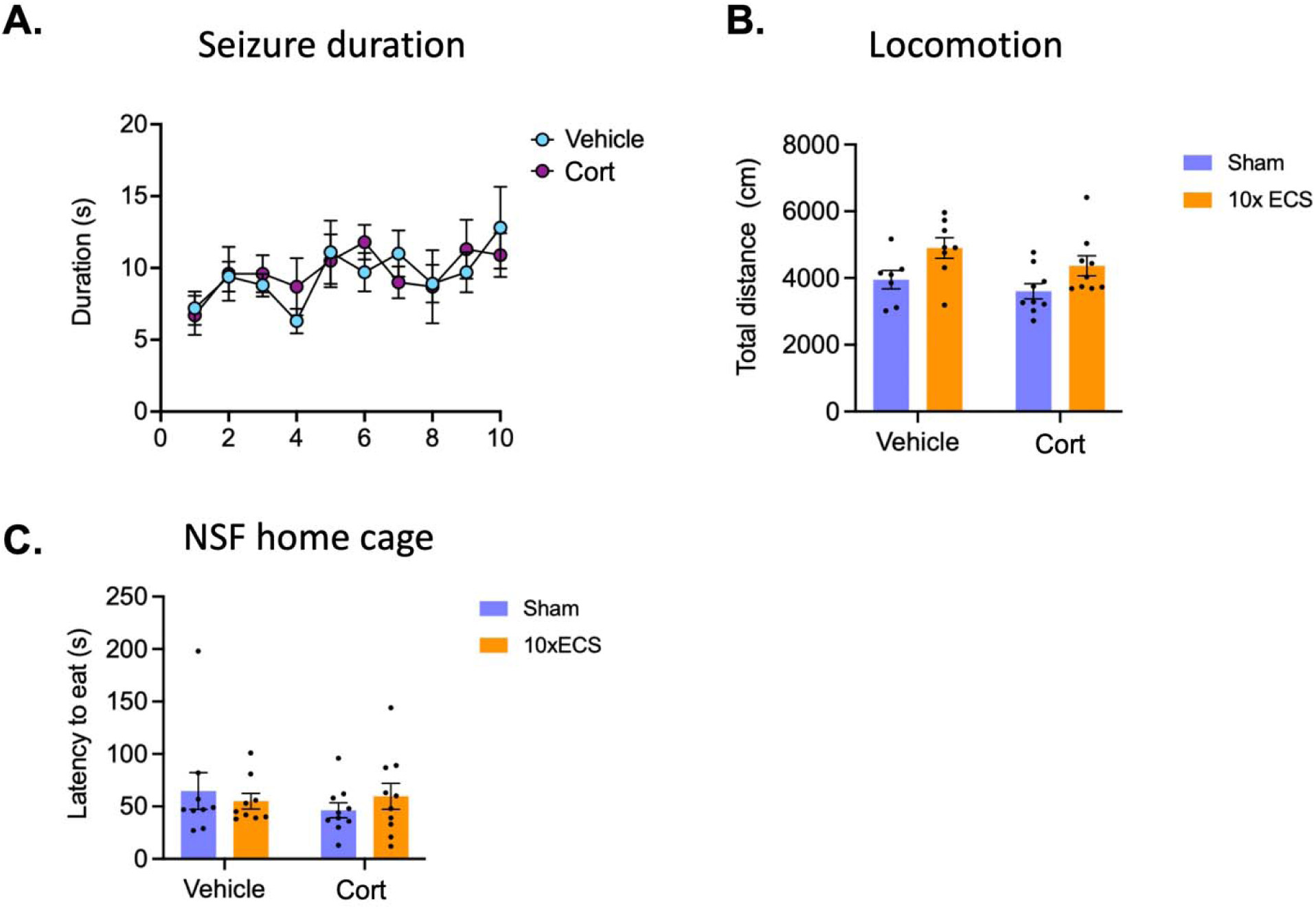
Additional characterization of the mouse model of ECS. A. Treatment with corticosterone (Cort, purple) vs vehicle (blue) had no impact on duration of seizure. B. Locomotion was measured in an open arena. While two-way ANOVA revealed an effect of ECS treatment (F (1, 29) = 9.200; p = .0051), post hoc Sidak’s test revealed no significant differences in vehicle (p = .0601) or Cort treated subjects (p = .1054). C. For the NSF, no differences in latency to eat in the home cage were observed.

## References

1. Rush AJ, South C, Jha MK, Jain SB, Trivedi MH. What to Expect When Switching to a Second Antidepressant Medication Following an Ineffective Initial SSRI: A Report From the Randomized Clinical STAR*D Study. J Clin Psychiatry 2020; 81(5).

2. Al-Harbi KS. Treatment-resistant depression: therapeutic trends, challenges, and future directions. Patient Prefer Adherence 2012; 6: 369–388.

3. Zhdanava M, Pilon D, Ghelerter I, Chow W, Joshi K, Lefebvre P et al. The Prevalence and National Burden of Treatment-Resistant Depression and Major Depressive Disorder in the United States. J Clin Psychiatry 2021; 82(2).

4. Husain MM, Rush AJ, Fink M, Knapp R, Petrides G, Rummans T et al. Speed of response and remission in major depressive disorder with acute electroconvulsive therapy (ECT): a Consortium for Research in ECT (CORE) report. J Clin Psychiatry 2004; 65(4): 485–491.

5. Ren J, Li H, Palaniyappan L, Liu H, Wang J, Li C et al. Repetitive transcranial magnetic stimulation versus electroconvulsive therapy for major depression: a systematic review and meta-analysis. Prog Neuropsychopharmacol Biol Psychiatry 2014; 51: 181–189.

6. Morcos N, Strominger J. Electroconvulsive Therapy for Unipolar Depression in Older Adults: Overall Outcomes and Clinical Trajectories of Nonresponders. J ect 2022; 38(4): 224–229.

7. Pagnin D, de Queiroz V, Pini S, Cassano GB. Efficacy of ECT in depression: a meta-analytic review. J ect 2004; 20(1): 13–20.

8. Santarelli L, Saxe M, Gross C, Surget A, Battaglia F, Dulawa S, et al. Requirement of hippocampal neurogenesis for the behavioral effects of antidepressants. Science 2003; 301(5634): 805–809.

9. Segi-Nishida E, Warner-Schmidt JL, Duman RS. Electroconvulsive seizure and VEGF increase the proliferation of neural stem-like cells in rat hippocampus. Proc Natl Acad Sci U S A 2008; 105(32): 11352–11357.

10. Ueno M, Sugimoto M, Ohtsubo K, Sakai N, Endo A, Shikano K et al. The effect of electroconvulsive seizure on survival, neuronal differentiation, and expression of the maturation marker in the adult mouse hippocampus. J Neurochem 2019; 149(4): 488–498.

11. Zhao C, Warner-Schmidt J, Duman RS, Gage FH. Electroconvulsive seizure promotes spine maturation in newborn dentate granule cells in adult rat. Dev Neurobiol 2012; 72(6): 937–942.

12. Schloesser RJ, Orvoen S, Jimenez DV, Hardy NF, Maynard KR, Sukumar M et al. Antidepressant-like Effects of Electroconvulsive Seizures Require Adult Neurogenesis in a Neuroendocrine Model of Depression. Brain Stimul 2015; 8(5): 862–867.

13. Meyers KT, Damphousse CC, Ozols AB, Campbell JM, Newbern JM, Hu C et al. Serial electroconvulsive Seizure alters dendritic complexity and promotes cellular proliferation in the mouse dentate gyrus; a role for Egr3. Brain Stimul 2023; 16(3): 889–900.

14. Nuninga JO, Mandl RCW, Boks MP, Bakker S, Somers M, Heringa SM et al. Volume increase in the dentate gyrus after electroconvulsive therapy in depressed patients as measured with 7T. Mol Psychiatry 2020; 25(7): 1559–1568.

15. Loef D, Tendolkar I, van Eijndhoven PFP, Hoozemans JJM, Oudega ML, Rozemuller AJM et al. Electroconvulsive therapy is associated with increased immunoreactivity of neuroplasticity markers in the hippocampus of depressed patients. Transl Psychiatry 2023; 13(1): 355.

16. Danielson NB, Kaifosh P, Zaremba JD, Lovett-Barron M, Tsai J, Denny CA et al. Distinct Contribution of Adult-Born Hippocampal Granule Cells to Context Encoding. Neuron 2016; 90(1): 101–112.

17. Drew LJ, Kheirbek MA, Luna VM, Denny CA, Cloidt MA, Wu MV et al. Activation of local inhibitory circuits in the dentate gyrus by adult-born neurons. Hippocampus 2016; 26(6): 763–778.

18. Li H, Tamura R, Hayashi D, Asai H, Koga J, Ando S et al. Silencing dentate newborn neurons alters excitatory/inhibitory balance and impairs behavioral inhibition and flexibility. Sci Adv 2024; 10(2): eadk4741.

19. McHugh SB, Lopes-Dos-Santos V, Gava GP, Hartwich K, Tam SKE, Bannerman DM et al. Adult-born dentate granule cells promote hippocampal population sparsity. Nat Neurosci 2022; 25(11): 1481–1491.

20. Anacker C, Luna VM, Stevens GS, Millette A, Shores R, Jimenez JC et al. Hippocampal neurogenesis confers stress resilience by inhibiting the ventral dentate gyrus. Nature 2018; 559(7712): 98–102.

21. Ash AM, Regele-Blasco E, Seib DR, Chahley E, Skelton PD, Luikart BW et al. Adult-born neurons inhibit developmentally-born neurons during spatial learning. Neurobiol Learn Mem 2023; 198: 107710.

22. Ikrar T, Guo N, He K, Besnard A, Levinson S, Hill A et al. Adult neurogenesis modifies excitability of the dentate gyrus. Front Neural Circuits 2013; 7: 204.

23. Trinchero MF, Giacomini D, Schinder AF. Dynamic interplay between GABAergic networks and developing neurons in the adult hippocampus. Curr Opin Neurobiol 2021; 69: 124–130.

24. Temprana SG, Mongiat LA, Yang SM, Trinchero MF, Alvarez DD, Kropff E et al. Delayed coupling to feedback inhibition during a critical period for the integration of adult-born granule cells. Neuron 2015; 85(1): 116–130.

25. Luna VM, Anacker C, Burghardt NS, Khandaker H, Andreu V, Millette A et al. Adult-born hippocampal neurons bidirectionally modulate entorhinal inputs into the dentate gyrus. Science 2019; 364(6440): 578–583.

26. Linden AM, Greene SJ, Bergeron M, Schoepp DD. Anxiolytic activity of the MGLU2/3 receptor agonist LY354740 on the elevated plus maze is associated with the suppression of stress-induced c-Fos in the hippocampus and increases in c-Fos induction in several other stress-sensitive brain regions. Neuropsychopharmacology 2004; 29(3): 502–513.

27. Munguba H, Gutzeit VA, Srivastava I, Kristt M, Singh A, Vijay A et al. Projection-Targeted Photopharmacology Reveals Distinct Anxiolytic Roles for Presynaptic mGluR2 in Prefrontal- and Insula-Amygdala Synapses. bioRxiv 2024.

28. Yoshimizu T, Shimazaki T, Ito A, Chaki S. An mGluR2/3 antagonist, MGS0039, exerts antidepressant and anxiolytic effects in behavioral models in rats. Psychopharmacology (Berl) 2006; 186(4): 587–593.

29. Maynard KR, Hobbs JW, Rajpurohit SK, Martinowich K. Electroconvulsive seizures influence dendritic spine morphology and BDNF expression in a neuroendocrine model of depression. Brain Stimul 2018; 11(4): 856–859.

30. Dranovsky A, Picchini AM, Moadel T, Sisti AC, Yamada A, Kimura S et al. Experience dictates stem cell fate in the adult hippocampus. Neuron 2011; 70(5): 908–923.

31. David DJ, Samuels BA, Rainer Q, Wang JW, Marsteller D, Mendez I et al. Neurogenesis-dependent and -independent effects of fluoxetine in an animal model of anxiety/depression. Neuron 2009; 62(4): 479–493.

32. Dieterich A, Srivastava P, Sharif A, Stech K, Floeder J, Yohn SE et al. Chronic corticosterone administration induces negative valence and impairs positive valence behaviors in mice. Transl Psychiatry 2019; 9(1): 337.

33. Bertholomey ML, Nagarajan V, Smith DM, Torregrossa MM. Sex- and age-dependent effects of chronic corticosterone exposure on depressive-like, anxiety-like, and fear-related behavior: Role of amygdala glutamate receptors in the rat. Front Behav Neurosci 2022; 16: 950000.

34. Wu MV, Hen R. Functional dissociation of adult-born neurons along the dorsoventral axis of the dentate gyrus. Hippocampus 2014; 24(7): 751–761.

35. Tuncdemir SN, Grosmark AD, Chung H, Luna VM, Lacefield CO, Losonczy A et al. Adult-born granule cells facilitate remapping of spatial and non-spatial representations in the dentate gyrus. Neuron 2023; 111(24): 4024–4039.e4027.

36. Hill AS, Sahay A, Hen R. Increasing Adult Hippocampal Neurogenesis is Sufficient to Reduce Anxiety and Depression-Like Behaviors. Neuropsychopharmacology 2015; 40(10): 2368–2378.

37. Nguyen PT, Dorman LC, Pan S, Vainchtein ID, Han RT, Nakao-Inoue H et al. Microglial Remodeling of the Extracellular Matrix Promotes Synapse Plasticity. Cell 2020; 182(2): 388–403.e315.

38. Butler A, Hoffman P, Smibert P, Papalexi E, Satija R. Integrating single-cell transcriptomic data across different conditions, technologies, and species. Nat Biotechnol 2018; 36(5): 411–420.

39. Satija R, Farrell JA, Gennert D, Schier AF, Regev A. Spatial reconstruction of single-cell gene expression data. Nat Biotechnol 2015; 33(5): 495–502.

40. Suzuki M, Masuda Y. Effect of repeated electroconvulsive shock treatment on a depression model, mouse forced swimming. Tohoku J Exp Med 1999; 189(1): 83–86.

41. Ittasakul P, Vora-Arporn S, Waleeprakhon P, Tor PC. Number of Electroconvulsive Therapy Sessions required for Thai Psychiatric Patients: a Retrospective Study. Neuropsychiatr Dis Treat 2020; 16: 673–679.

42. Henningsen K, Woldbye DP, Wiborg O. Electroconvulsive stimulation reverses anhedonia and cognitive impairments in rats exposed to chronic mild stress. Eur Neuropsychopharmacol 2013; 23(12): 1789–1794.

43. Samuels BA, Hen R. Novelty-Suppressed Feeding in the Mouse. Neuromethods 2011; 63: 14.

44. Kobayashi K, Ikeda Y, Sakai A, Yamasaki N, Haneda E, Miyakawa T et al. Reversal of hippocampal neuronal maturation by serotonergic antidepressants. Proc Natl Acad Sci U S A 2010; 107(18): 8434–8439.

45. Daval JL, Nakajima T, Gleiter CH, Post RM, Marangos PJ. Mouse brain c-fos mRNA distribution following a single electroconvulsive shock. J Neurochem 1989; 52(6): 1954–1957.

46. Brunner J, Ster J, Van-Weert S, Andrási T, Neubrandt M, Corti C et al. Selective silencing of individual dendritic branches by an mGlu2-activated potassium conductance in dentate gyrus granule cells. J Neurosci 2013; 33(17): 7285–7298.

47. Kamiya H, Ozawa S. Dual mechanism for presynaptic modulation by axonal metabotropic glutamate receptor at the mouse mossy fibre-CA3 synapse. J Physiol 1999; 518 ( Pt 2)(Pt 2): 497–506.

48. Ma J, Hu Z, Yue H, Luo Y, Wang C, Wu X et al. GRM2 Regulates Functional Integration of Adult-Born DGCs by Paradoxically Modulating MEK/ERK1/2 Pathway. J Neurosci 2023; 43(16): 2822–2836.

49. Gonzalez JC, Epps SA, Markwardt SJ, Wadiche JI, Overstreet-Wadiche L. Constitutive and Synaptic Activation of GIRK Channels Differentiates Mature and Newborn Dentate Granule Cells. J Neurosci 2018; 38(29): 6513–6526.

50. Samuels BA, Anacker C, Hu A, Levinstein MR, Pickenhagen A, Tsetsenis T et al. 5-HT1A receptors on mature dentate gyrus granule cells are critical for the antidepressant response. Nat Neurosci 2015; 18(11): 1606–1616.

51. Koumoundourou A, Rannap M, De Bruyckere E, Nestel S, Reissner C, Egorov AV et al. Regulation of hippocampal mossy fiber-CA3 synapse function by a Bcl11b/C1ql2/Nrxn3(25b+) pathway. Elife 2024; 12.

52. Mahar I, MacIsaac A, Kim JJ, Qiang C, Davoli MA, Turecki G et al. Effects of neuregulin-1 administration on neurogenesis in the adult mouse hippocampus, and characterization of immature neurons along the septotemporal axis. Sci Rep 2016; 6: 30467.

53. Ortega-Gascó A, Parcerisas A, Hino K, Herranz-Pérez V, Ulloa F, Elias-Tersa A et al. Regulation of young-adult neurogenesis and neuronal differentiation by neural cell adhesion molecule 2 (NCAM2). Cereb Cortex 2023; 33(21): 10931–10948.

54. Hagihara H, Ohira K, Miyakawa T. Transcriptomic evidence for immaturity induced by antidepressant fluoxetine in the hippocampus and prefrontal cortex. Neuropsychopharmacol Rep 2019; 39(2): 78–89.

55. Mendez-David I, David DJ, Deloménie C, Tritschler L, Beaulieu JM, Colle R et al. A complex relation between levels of adult hippocampal neurogenesis and expression of the immature neuron marker doublecortin. Hippocampus 2023; 33(10): 1075–1093.

56. Tosoni G, Ayyildiz D, Bryois J, Macnair W, Fitzsimons CP, Lucassen PJ et al. Mapping human adult hippocampal neurogenesis with single-cell transcriptomics: Reconciling controversy or fueling the debate? Neuron 2023; 111(11): 1714–1731.e1713.

57. Hochgerner H, Zeisel A, Lönnerberg P, Linnarsson S. Conserved properties of dentate gyrus neurogenesis across postnatal development revealed by single-cell RNA sequencing. Nat Neurosci 2018; 21(2): 290–299.

